# Model-based inference of a plant-specific dual role for HOPS in regulating guard cell vacuole fusion

**DOI:** 10.1101/2023.11.07.565947

**Authors:** Charles Hodgens, DT Flaherty, Anne-Marie Pullen, Imran Khan, Nolan J English, Lydia Gillan, Marcela Rojas-Pierce, Belinda S Akpa

## Abstract

Stomata are the pores on a leaf surface that regulate gas exchange. Each stoma consists of two guard cells whose movements regulate pore opening and thereby control CO_2_ fixation and water loss. Guard cell movements depend in part on the remodeling of vacuoles, which have been observed to change from a highly fragmented state to a fused morphology during stomata opening. This change in morphology requires a membrane fusion mechanism that responds rapidly to environmental signals, allowing plants to respond to diurnal and stress cues. With guard cell vacuoles being both large and responsive to external signals, stomata represent a unique system in which to delineate mechanisms of membrane fusion.

Fusion of vacuole membranes is a highly conserved process in eukaryotes, with key roles played by two multi-subunit complexes: HOPS (homotypic fusion and vacuolar protein sorting) and SNARE (soluble NSF attachment protein receptor). HOPS is a vacuole tethering factor that is thought to chaperone SNAREs from apposing vacuole membranes into a fusion-competent complex capable of rearranging membranes. To resolve a counter-intuitive observation regarding the role of HOPS in regulating plant vacuole morphology, we derived a quantitative model of vacuole fusion dynamics and used it to generate testable predictions about HOPS-SNARE interactions. We derived our model by applying simulation-based inference to integrate prior knowledge about molecular interactions with limited, qualitative observations of emergent vacuole phenotypes. By constraining the model parameters to yield the emergent outcomes observed for stoma opening – as induced by two distinct chemical treatments – we predicted a dual role for HOPS and identified a stalled form of the SNARE complex that differs from phenomena reported in yeast. We predict that HOPS has contradictory actions at different points in the fusion signaling pathway, promoting the formation of SNARE complexes, but limiting their activity.

**Author summary:** Plants “breathe” through pores in their leaves where each pore is formed by two specialized cells called guard cells. To open these pores, guard cells change in volume. This volume change is controlled by water-filled organelles called vacuoles that morph from multiple small entities to a few large ones capable of taking up more water to reshape the cell. Specialized proteins in vacuole membranes make this change happen by pulling vacuoles together until they fuse. Some of these proteins reside in membranes, but others must be drawn to the membrane from the cell’s cytoplasm. Specific lipid molecules in the membrane play an important role in recruiting those proteins to the vacuole membrane. We previously made an unexpected finding that removing this lipid induces plant vacuole fusion. To make sense of this observation, we used a mathematical model to piece together our knowledge of the proteins involved in this process and what we know about the chemical treatments that cause vacuoles to morph. Using computer simulations, we uncovered new rules about how molecules interact in membranes to accomplish the task of vacuole fusion in plants. We think the rules uncovered through mathematical modeling allow plants to respond quickly to environmental cues.

## Introduction

Stomata are pores in the surface of plant leaves that are critical for gas exchange – as required for photosynthesis and the control of leaf transpiration and temperature. Stomatal movement (opening and closing) is tightly controlled in response to exogenous cues such as changes in light or temperature, and endogenous cues such as circadian regulation or hormone signaling [1]. The transition from closed to open stomata is a complex process with several well-studied component phenomena, including the activation of blue-light photoreceptors, K+ ion influx, and water uptake [2]. Studies by confocal and electron microscopy have shown dynamic changes in the morphology of the vacuole between closed and open stomata [3,4]. Specifically, in the closed state of the stoma, the vacuoles within guard cells exhibit a fragmented or highly convoluted morphology, appearing sometimes as numerous small organelles [3–6]. When the pore is open, these same cells exhibit a vacuole morphology typical of other mature plant cells – namely a single large vacuole, or few vacuoles, occupying most of the intracellular space. Importantly, vacuole membrane fusion is necessary for full opening of the pore [6]. This dynamic vacuole activity is not observed in most mature plant tissue, making the guard cells a unique model to study vacuole fusion.

Fusion of vacuole membranes is a highly conserved process in eukaryotes and is best described in yeast [7,8]. Two multi-subunit protein complexes act in concert to induce vacuole fusion, the homotypic fusion and vacuolar protein sorting (HOPS) and soluble NSF attachment protein receptor (SNARE). HOPS is a tethering complex that is recruited from the cytosol by active RAB proteins and the presence of specific phosphoinositides at the vacuole membrane [8]. HOPS then is thought to provide binding sites for SNAREs from apposing membranes and thereby promote the formation of gap-spanning *trans*-SNARE complexes to support fusion. HOPS was also proposed to proofread the fidelity of the *trans*-SNARE complex and protect it from disassembly [9,10]. In the case of plant vacuoles, RAB7 has been implicated in homotypic vacuole fusion upstream of HOPS recruitment [11], which also requires the accumulation of phosphatidylinositol 3-phosphate (PI3P) [12].

Thus, the series of events leading to plant vacuole fusion, an essential transformation for full opening of the stomata, would seem to be: (1) HOPS subunits arrive at the vacuole membrane, mediated, in part, by the presence of PI3P; (2) HOPS tethers a pair of vacuoles and chaperones their SNARE proteins into the *trans*-SNARE fusion machinery; (3) the *trans-*SNARE complex zippers, exerting the force required to fuse apposing membranes, an event that is accompanied by HOPS release from the membrane (Figure 1A). Linear logic would lead one to expect that withdrawing PI3P from this system would impair the cell’s ability to respond to pore-opening signals by fusing vacuoles. However, depleting PI3P by treating guard cells with the Phosphatidyl-Inositol 3-Kinase (PI3K) inhibitor wortmannin causes spontaneous fusion in plants (Figure 1B). That is, even in the absence of the appropriate environmental or biological cue, guard cell vacuoles fuse when PI3P, a membrane lipid thought to be required for forming the fusion machinery, is removed [6]. This seems to be a plant-specific process, as PI3P depletion does not induce vacuole or lysosome fusion in yeast or animal cells [13–15].

**Fig 1.**
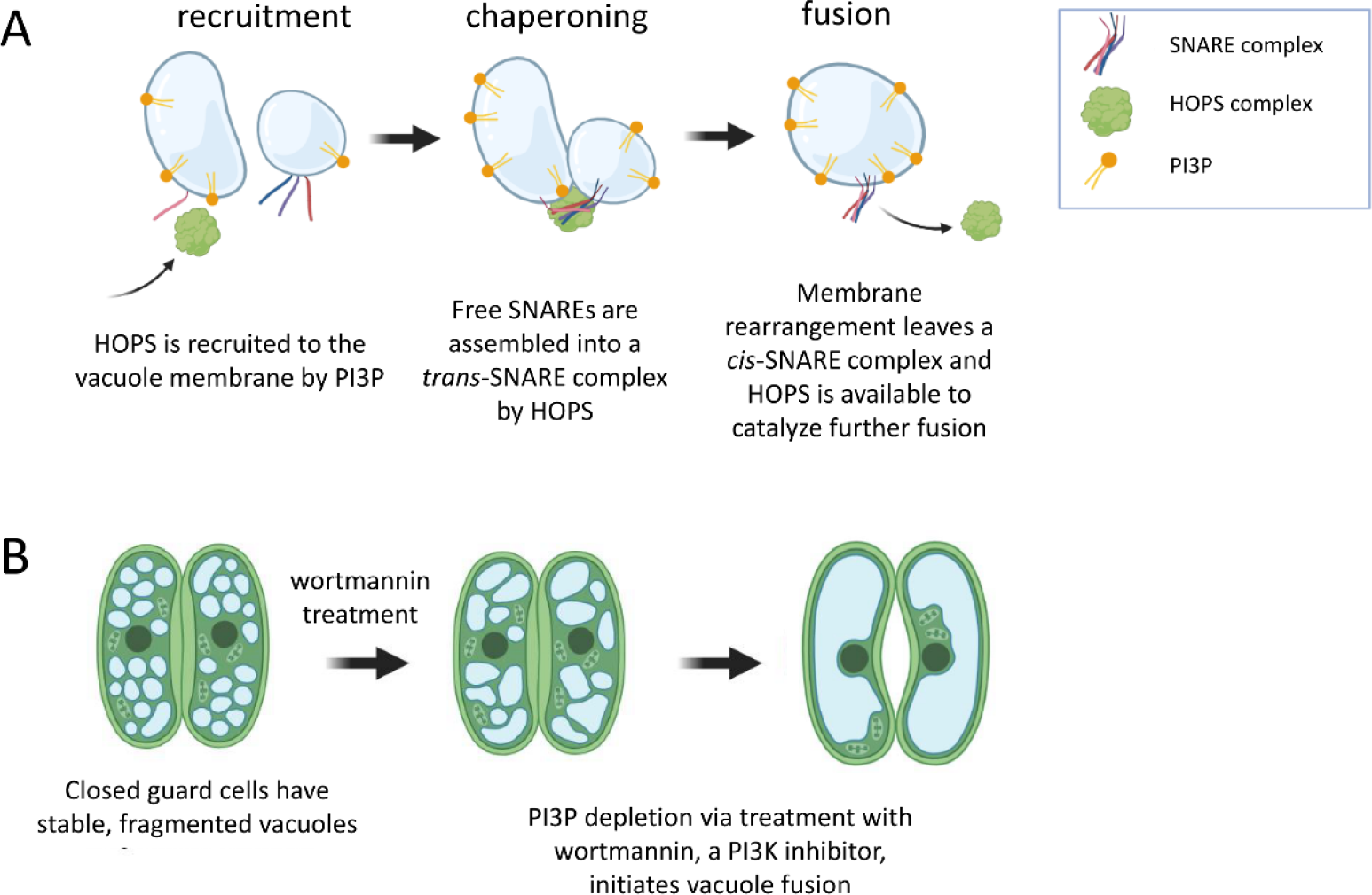
Contradictory results regarding the mode of action of PI3P and HOPS in plant vacuole fusion. (A) HOPS is recruited from a cytosolic pool to the vacuole membrane, due, in part, to the presence of the phosphoinositide PI3P. HOPS is responsible for tethering vacuoles together and chaperoning membrane-embedded SNARE proteins into a *trans*-SNARE fusion complex. Vacuole fusion is mediated by the zippering activity of the SNARE complex. As part of the fusion event, HOPS leaves the membrane. Based on this prior knowledge, one would expect fusion to be impossible in the absence of PI3P. (B) When wortmannin, a PI3K inhibitor, is used to deplete PI3P from the guard cells of closed stoma, small vacuoles rapidly fuse.

These observations present a puzzle as to the preconditions for vacuole fusion. To explore how PI3P depletion could promote spontaneous fusion, we developed a systems model of known and hypothetical events that may control fusion complex assembly and activation in *Arabidopsis*. Our model consists of ordinary differential equations (ODEs) describing the dynamic recruitment, complexation, and interaction of HOPS and SNARE proteins at vacuolar membranes. We used mass action kinetics to capture the dependence of each event on the abundance of required species. As with all such mathematical models, we then had the challenge of assigning values to rate constants and other parameters of the model. As directly measuring the kinetics of individual molecular-scale events is challenging, we sought to inform the kinetics of molecular events by simulation-based inference [16–21]. Essentially, knowing what system perturbations should promote fusion, and having some sense of how fusion rates differ under different perturbations, we can classify candidate parameter sets as plausible or implausible based on the model’s ability to predict the expected fusion dynamics. This means that we make no pretense of defining a single parameterization, but instead computationally pre-screen the model to exclude kinetics that are inconsistent with current knowledge and define the domain of kinetics that is worthy of further interrogation.

While computational models can offer a machine-assisted approach to reason about biological data, one typically desires abundant, quantitative data to inform such models. However, biological data often come in the form reported in Figure 2, which captures emergent, qualitative vacuole phenotype. Here, we do not have a finely resolved time-course of stomatal dynamics or vacuole number. In principle, one could derive quantitative information about vacuole size and number from live-cell images. That would require (i) a sufficient number of images to obtain statistically sound estimates, (ii) automated image segmentation algorithms to alleviate the laborious tasks of counting and measuring vacuoles, (iii) images of sufficient quality for successful application of such algorithms, and (iv) adequate financial and skilled human resources to perform the larger number of experiments required. By contrast, a qualitative summary of vacuole fusion phenomena is straightforward to make, and we sought to determine whether this information might be adequate to constrain our definition of plausible biological mechanisms via a mathematical model. If so, such a modeling approach could pre-screen likely regulatory mechanisms and inform a targeted experimental strategy in which to invest greater time and resources.

**Fig 2.**
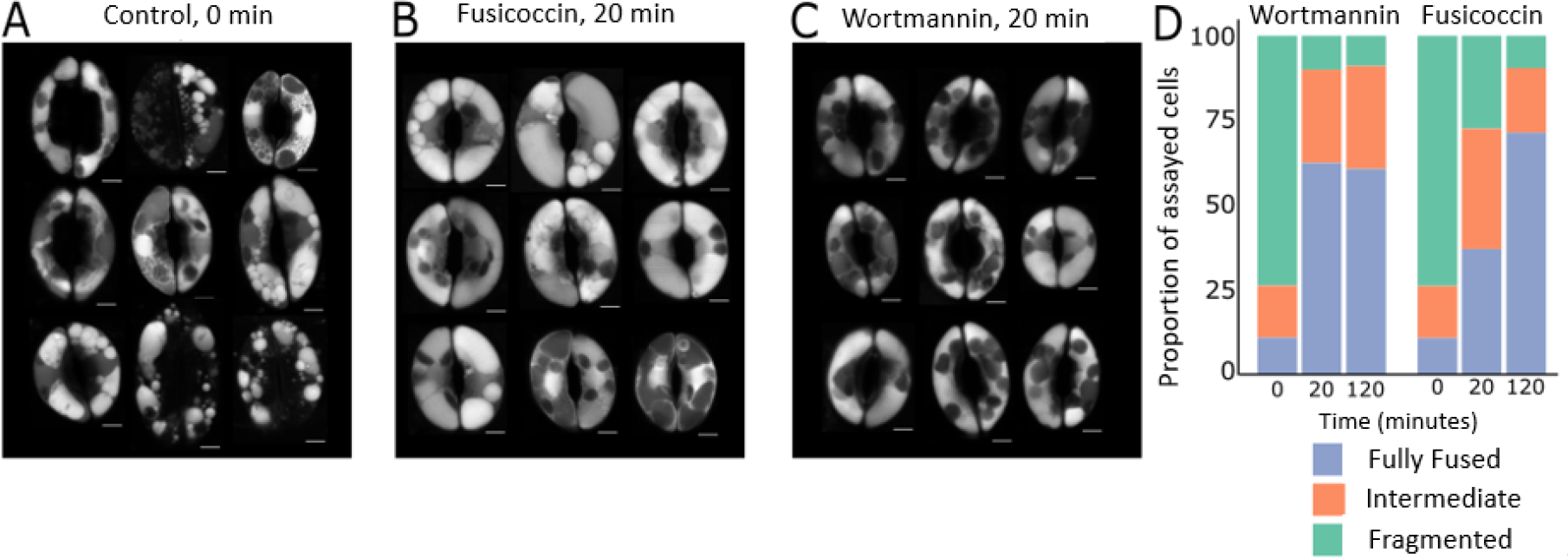
Fusion due to wortmannin treatment proceeds more quickly than fusion due to a fusicoccin stimulus. (A) Dark-acclimated, ABA-treated *Arabidopsis thaliana* guard cells were imaged prior to treatment with a fusion-inducing stimulus. (B) Vacuole morphology after 20 minutes of fusicoccin and light treatment and (C) after 20 minutes of wortmannin treatment. (D) A qualitative survey of vacuole morphology reveals rapid wortmannin-driven fusion. We classified guard cells as having fragmented, fully fused, or intermediate vacuole phenotypes at zero minutes, twenty minutes, and two hours after inducing fusion. The evolution of vacuole morphology was complete after 20 minutes in the wortmannin-treated cell group. Vacuole morphology in fusicoccin-treated cells continued to evolve after that time. Vacuoles were stained with BCECF. Chloroplasts (dark ovals inside guard cells) typically do not take up the vacuole stain.

## Results and Discussion

### PI3P depletion causes vacuoles to fuse faster than fusicoccin treatment does

Stoma vacuole fusion can be induced under laboratory conditions by treatment with fusicoccin or wortmannin. Fusicoccin is a fungal toxin that promotes stomata opening by activation of the plasma membrane H^+^ ATPase [22,23]. Thus, fusicoccin can be used as a proxy for the signaling pathway that triggers stomata opening downstream of light perception [23], but fusicoccin is not expected to drive vacuole fusion directly. The modeling strategy we report herein was motivated by experiments comparing the dynamics of fusicoccin- and wortmannin-induced vacuole fusion. We first induced stomata closure by incubating leaves with abscisic acid (ABA) in the dark. Consistent with prior results [6,24], this resulted in guard cells with highly fragmented vacuoles (Figure 2A). The leaves were subsequently treated with either fusicoccin or wortmannin, and both treatments induced vacuole fusion (Figure 2B and C, respectively).

Vacuoles in wortmannin-treated guard cells fused rapidly, completing this change in morphology within 20 minutes (Figure 2D, left). At that 20-minute timepoint, vacuoles fusing in response to fusicoccin still presented an intermediate morphology, with fusion activity continuing for over an hour (Figure 2D, right). Thus, guard cell vacuole fusion presents non-heuristic emergent dynamics. Specifically, (1) wortmannin depletes PI3P from vacuole membranes, and thereby initiates a series of events previously thought to depend on the presence of that lipid; and (2) the fusion process initiated by wortmannin treatment is accelerated compared to that associated with normal physiological responses (mimicked here by fusicoccin treatment). With the goal of proposing a molecular pathway capable of explaining these observations, we used mathematical modeling as a tool to integrate these phenotypic observations with prior knowledge about the molecular machinery of fusion.

### Prior knowledge of molecular mechanisms fails to explain PI3P regulation of fusion

By collating information about membrane fusion in plants, yeast, and animal cells, we established the following as prior knowledge about the molecular events leading to vacuole fusion: (1) Membrane fusion requires a *trans*-SNARE complex [25]; (2) HOPS chaperones free SNARE proteins into a *trans*-SNARE complex [26,27]; (3) The HOPS subunit VPS41 (AT1G08190) is observed at vacuole membranes only in the presence of the membrane lipid PI3P [12]. Together, these facts align with the linear scheme shown in Figure 3A, where the process of fusion appears to emerge as a direct consequence of the presence of PI3P. The scheme implies that the absence of PI3P would prevent recruitment of HOPS subunits to the vacuole membrane and would thus prevent chaperoning of the *trans-*SNARE fusion complex. However, our experiments yielded the unexpected observation that PI3P depletion causes vacuoles to fuse. Given that linear reasoning about the molecular mechanisms underlying vacuole fusion failed to explain the reality of the biology, we sought an alternative mechanism that could capture the complex outcomes observed – and do so while remaining consistent with our prior knowledge about the molecular players in the system.

**Fig 3.**
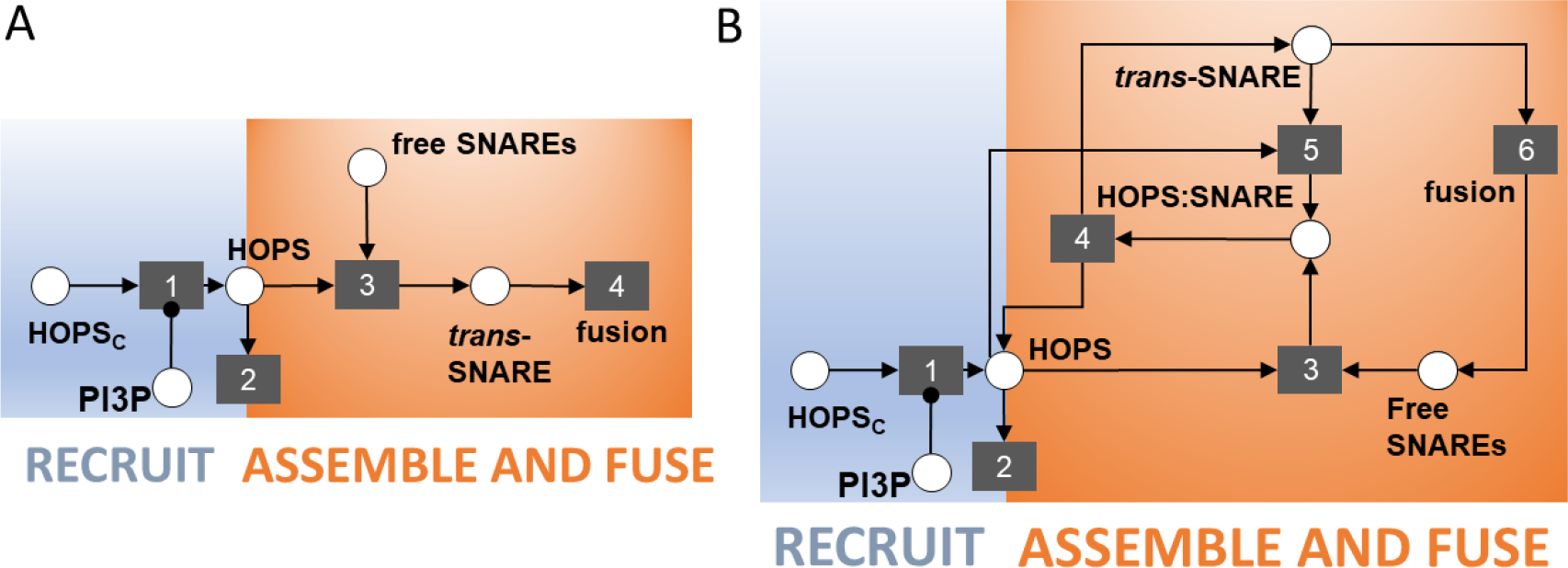
Two schematics of increasing complexity that describe potential molecular events leading to vacuole fusion. (A) A schematic of the known signaling events leading to vacuole fusion. Circles represent species and their complexes. Squares represent events. Arrows into an event indicate the logical requirements for that event to take place. Arrows originating from an event indicate the event’s consequences. The replacement of an arrowhead by a dot indicates that a species is required for an event to take place, but that species is not consumed when the event happens. Thus, PI3P is required for HOPS to be recruited from the cytosol to the membrane (event 1). HOPS may then leave the membrane (event 2) or participate in chaperoning of free SNARE proteins into a *trans*-SNARE complex (event 3). Finally, the *trans*-SNARE complex drives fusion activity (event 4). This linear scheme and its cascading series of logical requirements implies that the removal of PI3P can only prevent fusion. (B) Schematic of increased complexity positing the spontaneous dissociation (and reassociation) of a HOPS:*trans*-SNARE super-complex. **Events 1 and 2 are retained from part A**. HOPS chaperoning of free SNARES (event 3) results in a HOPS:*trans*-SNARE super-complex from which HOPS must be removed – if we assume that the *trans*-SNARE complex is the competent species driving fusion (event 6). The simplest way we can posit for HOPS to be removed is by spontaneous dissociation (event 4). Species that dissociate spontaneously can likely reassociate (event 5).

We began by asking what additional complexity might have been overlooked in our simple, linear description of the biological mechanism. For one, if HOPS chaperones SNARE proteins into a *trans*-SNARE complex, then there is, at some point, a HOPS:*trans-*SNARE super-complex. This complex’s existence was previously implicit in the chaperoning event (event 3 in Figure 3A). Making it explicit in the model compels us to ask whether the HOPS:*trans*-SNARE super-complex is competent for facilitating fusion. The alternative would be to posit that the bare *trans*-SNARE complex is the fusion-competent assembly. We proceeded with this second assumption, which creates a requirement for HOPS to be removed from the super-complex. Following the principle of parsimony, we adopted the simplest possible explanation for HOPS removal – namely, spontaneous dissociation (Figure 3B, event 4). If two species can spontaneously dissociate, it is reasonable to consider the possibility of reassociation (Figure 3B, event 5). Finally, we made explicit the events that follow membrane fusion. After two membranes join, the SNARE complex has all members co-located in the same membrane, as a *cis*-SNARE complex. This must be disassembled to prime free SNAREs to participate in new vacuole-bridging complexes that drive subsequent rounds of vacuole fusion. We combined the events of fusion, disassembly, and priming into a single abstracted event 6, as shown in Figure 3B.

This non-linear scheme offers a potential explanation for the fusion response observed in the wortmannin experiment. If the fusion-competent species is the bare *trans*-SNARE complex, once the HOPS:*trans*-SNARE super-complex is formed, any perturbation that promotes the dissociated state will lead to fusion activity. If dissociation of the super-complex were reversible, an excess of HOPS in the membrane would tend to keep the *trans-*SNARE complex in the super-complex state. Conversely, any perturbation that reduces HOPS abundance in the membrane would promote accumulation of the fusion-competent *trans*-SNARE machinery. In this schematic, two competing processes determine HOPS abundance in the membrane: HOPS subunit recruitment (event 1), which requires PI3P, and HOPS turnover from the membrane (event 2), which we assume to proceed at some basal rate. Depleting PI3P would put a stop to HOPS recruitment, leaving the turnover process to eliminate HOPS from the vacuole membrane. If turnover were sufficiently rapid, the result would be a swift release of fusion competent *trans*-SNARE complexes and corresponding fusion activity.

While this conceptual model may explain the wortmannin experiment, it does not allow for fusion in the presence of PI3P. As PI3P depletion is a consequence of a lab-based chemical perturbation and is not known to be a feature of vacuole fusion in the native plant, the model is, at best, incomplete. However, the framework defines a hypothetical function required to complete the formation of fusion-competent SNARE complexes – one that enables a search for a missing signal in our biological mechanism.

### Conceptual model suggests a missing signaling event, and yeast vacuole fusion offers a candidate signal

Our conceptual model suggests that HOPS removal could be a critical event to activate the *trans*-SNARE complex. Thus, we posited that the native stoma might possess an active mechanism for HOPS displacement from fusion complexes. While absent from the *Arabidopsis* literature, we found evidence of such stalling and activation phenomena amongst the comparatively well-studied proteins involved in yeast vacuole fusion. Specifically, we noted the example of Sec17 (SGD:S000000146), an alpha NSF attachment protein (α-SNAP) with a multifaceted role in yeast membrane fusion [28– 33]. Sec17 facilitates rapid fusion when added to mixtures of stalled intermediate complexes in reconstituted proteoliposome experiments [33,34] involving truncated SNARE proteins which cannot fully zipper together to drive membrane fusion. Interactions between membrane lipids and an apolar loop on Sec17 lower the energy barrier for membrane rearrangement, thereby encouraging fusion [33].

Given this information, we asked whether *Arabidopsis* possesses homologs of yeast Sec17. A BLASTP search of the *Arabidopsis* genome returns two loci with high similarity to Sec17: one locus with two isoforms (AT3G56190, E-values of 3e-38 and 1e-29) and another with one form (AT3G56450, E-value of 5e-15). These loci are annotated as ASNAP/ALPHA-SOLUBLE NSF ATTACHMENT PROTEIN (SNAP)2 (AT3G56190) and ALPHA-SNAP1 (AT3G56450). Roles for these proteins in the *Arabidopsis* fusion apparatus have not been reported, but a role in gametogenesis for ASNAP/ALPHA-SNAP2 has been established by Liu *et al*. [35]. Finally, both ASNAP1 and ASNAP2 transcripts are present in *Arabidopsis* guard cells [36,37]. Thus, we adopted Sec17 as a candidate signal for displacement of HOPS from HOPS-*trans*SNARE super-complexes, as depicted in the updated model shown in Figure 4. However, we wish to reiterate our hypothesis concerns a functional role, not a specific protein. While these proteins are strong candidates, it may be the case that another protein besides ASNAP1 or ASNAP2 performs this function in *Arabidopsis*. Notwithstanding, the schematic in Figure 4 informs the remainder of this investigation.

**Fig 4.**
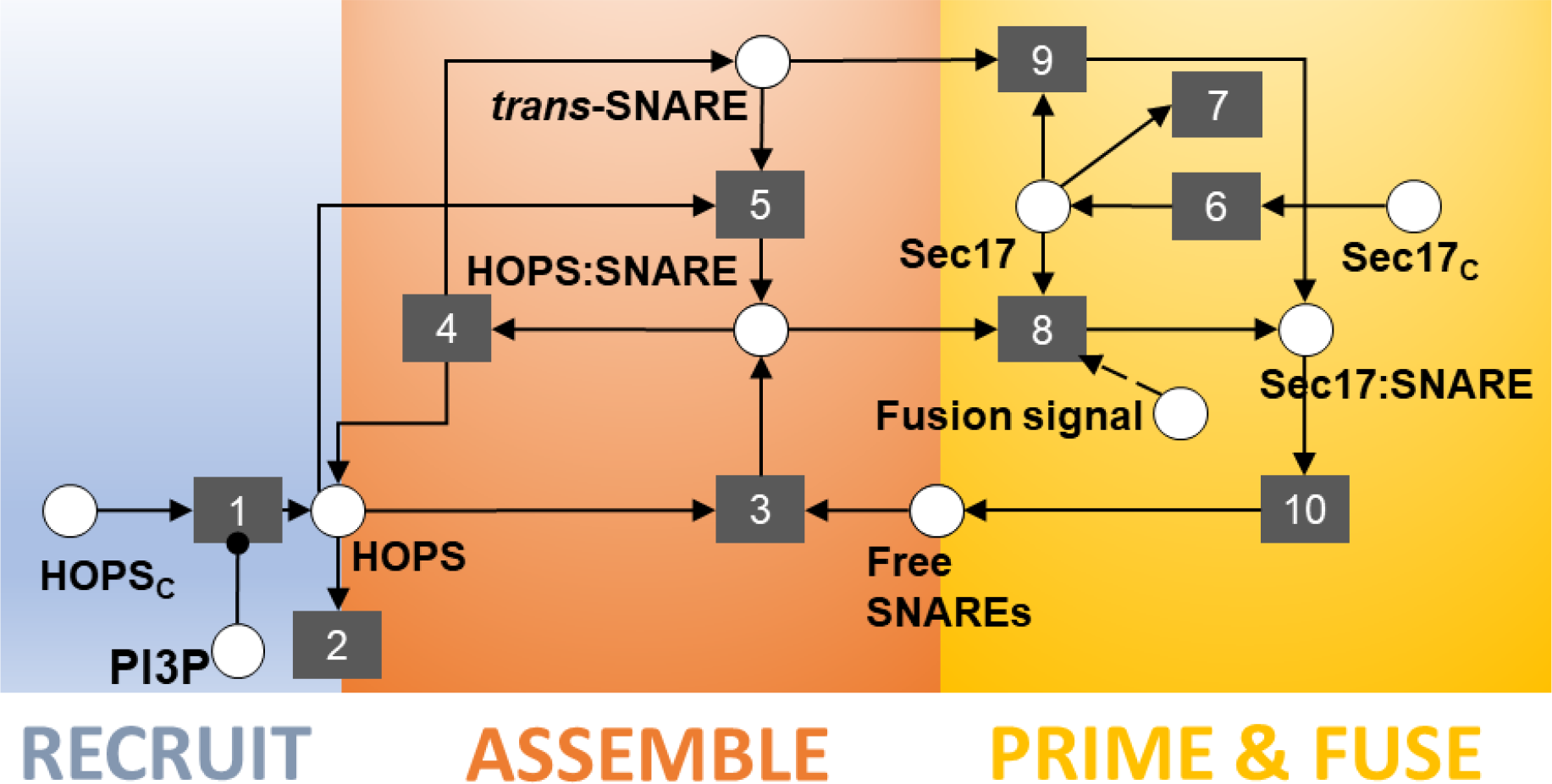
Final schematic offering the hypothesis that HOPS displacement is required to activate the *trans*-SNARE complex and drive fusion. The literature on yeast vacuole fusion offers a candidate signal with precisely that function – *i*.*e*., activating assembled, but inactive, trans-SNARE complexes. Furthermore, the *Arabidopsis* analogue of that yeast protein is expressed in guard cells. In our schematic, this protein, Sec17, is a cytoplasmic species recruited to the membrane (event 6), where it can associate with and activate *trans*-SNARE complexes (event 9). We posit that Sec17 cannot displace HOPS from HOPS:*trans*-SNARE super-complexes until the system receives some upstream signal that triggers stoma opening (event 8). After HOPS displacement, the fusion-competent complex drives fusion events (event 10). We allow the abstraction in event 10 to be inclusive of post-fusion phenomena such as disassembly of the *cis*-SNARE complex, freeing individual SNARE proteins to participate in further rounds of fusion. Event 7 represents turnover of Sec17 from the membrane. Events 1 through 5 are retained from earlier schematics (Figure 3).

Determining the validity of this conceptual model required that we confront it with empirical data. To this end, we turned the diagram in Figure 4 into a system of differential equations that we could simulate to predict emergent outcomes under different perturbations. The equations, reported in the detailed methods section, reflect the evolution of membrane protein abundances and configurations over time. When non-dimensionalized, the system of equations featured eight parameters. Table 1 defines those parameters, along with their associated events in the conceptual model.

**Table 1.** Model parameters and their descriptions.

| parameter | corresponding event | description |
| --- | --- | --- |
| $K_1$ | 1,2 | rates of HOPS recruitment and turnover |
| $K_3$ | 3 | chaperoning rate |
| $K_4$ | 4 | rate of HOPS:*trans*-SNARE super-complex dissociation |
| $\alpha (= K_5/K_4)$ | 5 | HOPS & *trans*-SNARE reassociation rate expressed relative to HOPS:*trans*-SNARE dissociation rate |
| $K_6$ | 6 | rate of Sec17 recruitment |
| $K_7$ | 7 | rate of Sec17 turnover |
| $\beta (= K_8/K_9)$ | 8 | rate for Sec17 displacement of HOPS expressed relative to rate for Sec17 association with *trans*-SNARE complexes |
| $K_9$ | 9 | rate of Sec17 association with *trans*-SNARE complexes |

The only data available to inform the model (Figure 2) reports on the evolution of vacuole morphology and does not capture the kinetics of individual events represented in the model. Assuming that the model parameters have been determined to be structurally identifiable, parameter inference approaches can be used to reverse engineer parameter values based on known systems-level outcomes. However, that typically involves quantitative observations, and, preferably, time-series data of sufficient quality and quantity to render parameters practically identifiable. With only qualitative phenotypic observations to work with, we questioned whether that information could meaningfully constrain the parameterization of our proposed model and whether we could learn anything about the properties of the molecular signaling pathway from the available data.

### Vacuole morphology, as a qualitative phenotype, constrains model parameterization

As a first attempt to assess whether the existing observations might meaningfully constrain model parameterization, we ran simulations using parameter values randomly sampled across a large range spanning eight orders of magnitude. While sampling ∼10^5^ different parameter sets could not even begin to meaningfully explore the eight-dimensional parameter space, this cursory assessment did shed some light on whether the expected fusion characteristics are trivial to produce. To this end, we specified semi-quantitative emergent behaviors that the simulations would need to produce for the predictions to match our biological observations. We defined a match as any simulation that met five criteria: (1) the system must have a stable steady state prior to any perturbation causing fusion activity; (2) the system should exhibit increased fusion activity upon removal of PI3P; (3) the system should exhibit increased fusion activity upon triggering of event 8, our hypothetical mechanism of fusicoccin-responsive fusion; (4) fusion due to PI3P removal should occur more quickly than fusion due to trigger activation; (5) spontaneous fusion events in the absence of a specific signaling perturbation should be rare. We converted these qualitative statements into quantitative criteria (CR_i_) by setting numerical thresholds (TH_i_) for acceptance of any given simulation. Table 2 details these acceptance criteria, expressed as inequalities.

**Table 2.**
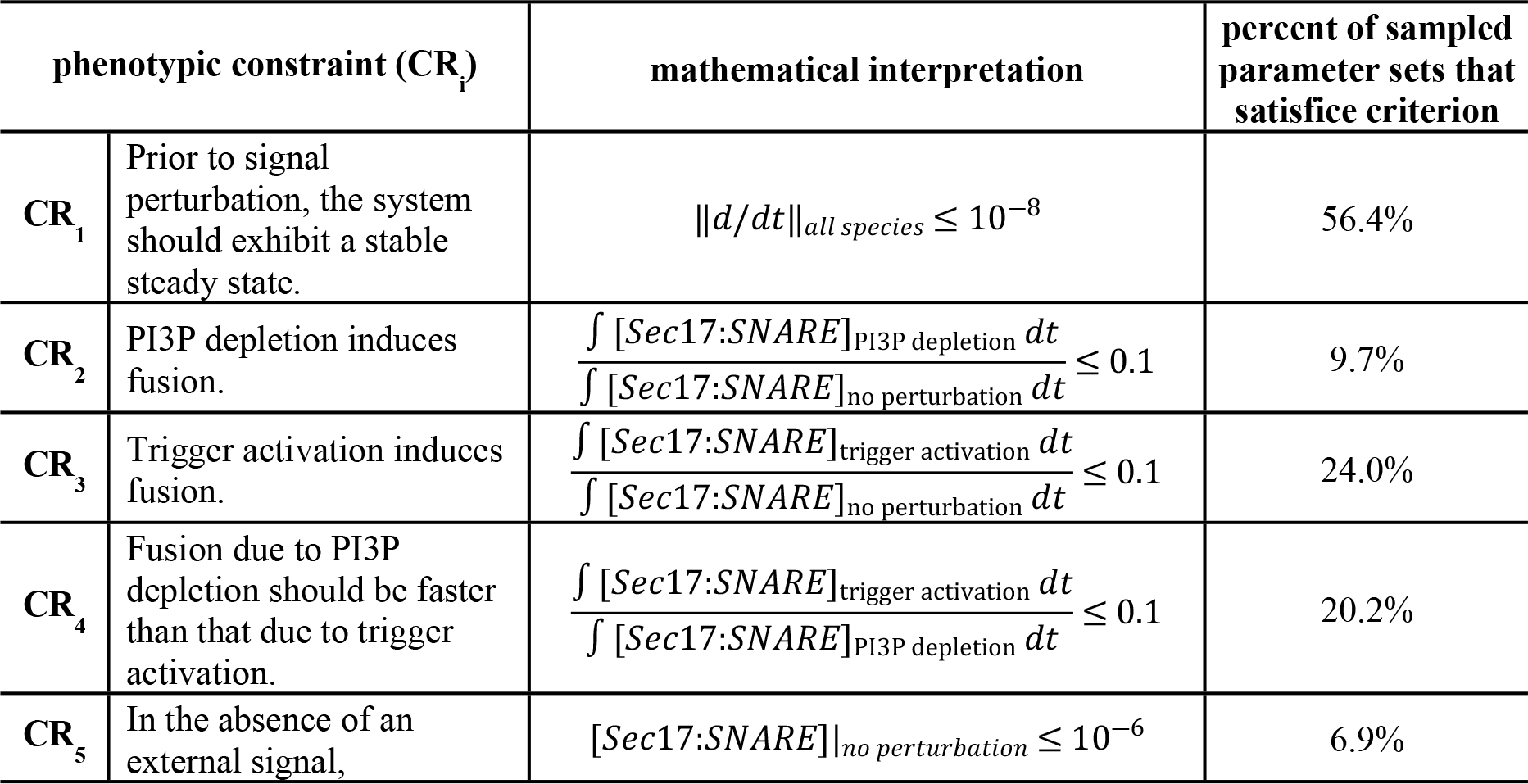
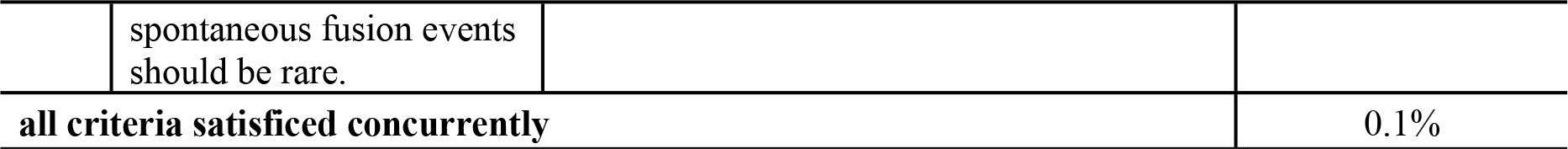
Translating qualitative phenotype observations into quantitative constraints.

| phenotypic constraint ($CR_i$) | | mathematical interpretation | percent of sampled parameter sets that satisfy criterion |
| --- | --- | --- | --- |
| $CR_1$ | Prior to signal perturbation, the system should exhibit a stable steady state. | $\ d/dt\ _{all\ species} \leq 10^{-8}$ | 56.4% |
| $CR_2$ | PI3P depletion induces fusion. | $\frac{\int [Sec17:SNARE]_{PI3P\ depletion} dt}{\int [Sec17:SNARE]_{no\ perturbation} dt} \leq 0.1$ | 9.7% |
| $CR_3$ | Trigger activation induces fusion. | $\frac{\int [Sec17:SNARE]_{trigger\ activation} dt}{\int [Sec17:SNARE]_{no\ perturbation} dt} \leq 0.1$ | 24.0% |
| $CR_4$ | Fusion due to PI3P depletion should be faster than that due to trigger activation. | $\frac{\int [Sec17:SNARE]_{trigger\ activation} dt}{\int [Sec17:SNARE]_{PI3P\ depletion} dt} \leq 0.1$ | 20.2% |
| $CR_5$ | In the absence of an external signal, | $[Sec17:SNARE]_{no\ perturbation} \leq 10^{-6}$ | 6.9% |

|  |  |  |
| --- | --- | --- |
|  | spontaneous fusion events<br>should be rare. |  |
| **all criteria satisfied concurrently** |  | 0.1% |

As our model can only report the instantaneous flux associated with the fusion event (*i*.*e*., a rate proportional to the abundance of [Sec17:SNARE], as indicated by event 10 in Figure 4), we chose to quantify cumulative fusion activity as the integral under the curve of fusion rate vs. time. For a given parameter set, we compared these integrals across simulations performed without a signal perturbation to those with a signal perturbation. As we had no quantitative data on relative fusion dynamics, we selected very conservative thresholds for the expected changes in fusion activity. For example, for criterion 2, when we simulated PI3P removal, we accepted any simulation increasing fusion activity by ten-fold or greater. Biological intuition suggests that a ten-fold increase is likely insufficient to explain our guard cell observations. If vacuole fusion takes place within 20 minutes (Figure 2), and that process involves fusion activity only ten-fold greater than that in untreated cells, one might expect to see vacuoles fuse spontaneously in untreated cells over a several hours. This does not occur, so our chosen threshold is likely overly permissive. However, an overly restrictive threshold might elide regions of parameter space that are, in truth, biologically relevant. We chose to err on the side of permissiveness.

Table 2 reports the mathematical definitions of our criteria and our chosen acceptance thresholds. We accepted as plausible any parameterization that produced simulation results that met these thresholds. In this Table, we also detail the fraction of the evaluated parameter sets that returned emergent dynamics matching this quantitative interpretation of our qualitative biological observations. These fractions were determined by evaluating model outcomes for N=2^18^ points in the eight-dimensional parameter space. We generated these samples using uniform Sobol’ sampling (*sobolset* function in MATLAB) and found that less than 1% of the examined parameter sets produced simulation outcomes matching all five of our desired emergent behaviors. As fully satisficing parameter sets were scarce, we posited that our conservative criteria, expressed as inequalities, may be sufficient to constrain parameterization of our model. The parameters would clearly not be uniquely identifiable, but we should be able to discriminate subdomains of parameter space that return biologically plausible kinetics. Interrogation of those sub-domains could then indicate experimentally tractable measurements that would permit us to establish the viability of our hypothesized signaling pathway. A simulation-based inference approach such as approximate Bayesian computation can enable a search for those domains by identifying plausible parameter values.

### Simulation-based inference delineates plausible regions of parameter space

As we approached the problem of parameterizing this model, we first determined whether the model might be insensitive to any of the eight parameters. If so, one could fix the value of one or more parameters and thereby simplify our search for plausible kinetic constants. To this end, we performed a Sobol’ global sensitivity analysis using the correlation-based approach of Glen and Isaacs [38]. This variance-based approach returns indices indicating how much each model parameter contributes to the variance of each simulation outcome. Performing the analysis required that we turn our simulation acceptance criteria into quantitative metrics indicating how far a given simulation is from satisficing the established criteria. We chose to translate each criterion and its acceptance threshold via a RelU objective function, as indicated in Equation 1. This formulation has the benefit of mapping all results meeting the acceptance threshold to identical scores of zero, while penalizing results that do not meet our thresholds. Failing simulations will return scores that increase linearly as the simulation outcomes increasingly deviate from our desired behaviors. Using these functions as the outcomes evaluated in the Sobol’ analysis allowed us to evaluate how each model parameter contributed to a decision as to whether a simulation would produce satisficing emergent behaviors.

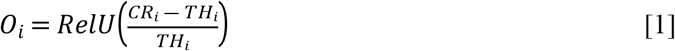

Figure 5 reports our estimates of the first order and total sensitivity indices. We note that acceptance of the steady state criterion appears insensitive to all parameters when only first-order effects are considered. However, when second and higher order parameter interactions are included (*i*.*e*., in total sensitivity index), the steady-state requirement exhibits the greatest sensitivity to all parameters. Although the parameter describing the rate constant for Sec17 displacement of HOPS contributes little variance to satisfaction of our five outcomes of interest, the parameter’s impacts on outcomes 3 and 4 (relative rates of fusion under different treatments and expectations for basal fusion, respectively) are significantly different from zero, as determined by a Wilcoxon Rank Sum test (T_i_ = 0.009 and 0.03, respectively; p < 1e-6. Approach detailed in Methods). Note that these values do not reflect absolute sensitivity, but rather indicate each parameter’s relative contribution to overall variance of each model outcome. Given the results of this sensitivity analysis, we chose to estimate all eight parameters.

**Fig 5.**
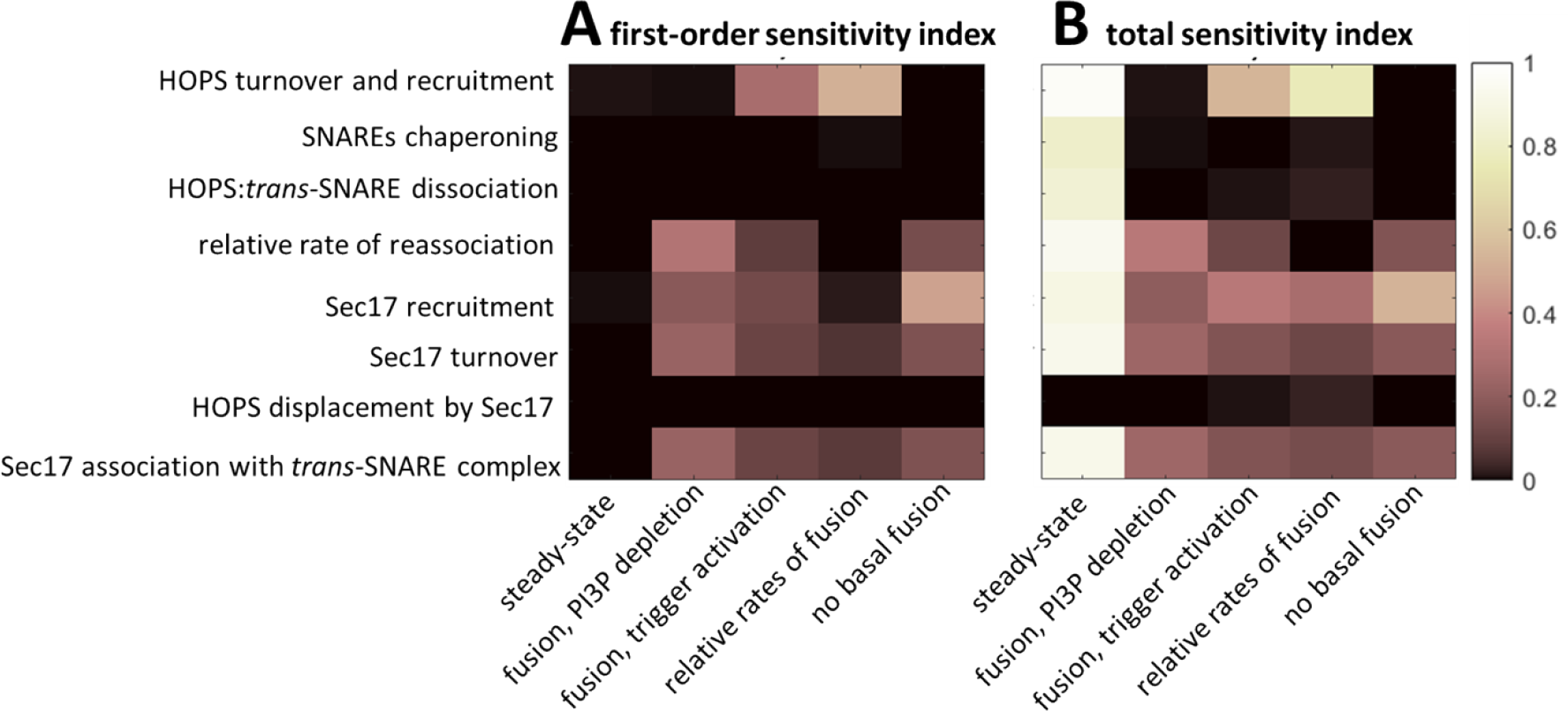
Variance-based global sensitivity analysis. (A) First-order indices, indicating what fraction each model parameter (y labels) independently contributes to the variance of each model outcome (x labels). (B) Total Sobol’ indices indicating each parameter’s contribution to the variance of each outcome when considering all inter-parameter interactions. We calculated the Sobol’ indices using two independent sets of N=10^5^ samples. Results indicate that all parameters make a statistically significant contribution to satisficing the desired model outcomes. However, the contribution of the rate constant for HOPS displacement by Sec17 is weak across all outcomes, and the SNARE chaperoning rate has little impact on fusion dynamics. Seven of the eight parameters strongly impact the steady state criterion, but they do so almost exclusively through parameter interactions.

Using our objective functions as indicators of valid emergent dynamics, we attempted to infer plausible regions of parameter space for our model. To achieve this, we used the Bayesian inference approach described by Toni *et al*. [39]. – namely, Approximate Bayesian Computation using Sequential Monte Carlo (ABC-SMC). This algorithm requires a distance metric to quantify the deviation of a simulation outcome from a target value. Given our desire to satisfice multiple outcomes simultaneously, we summed the five objective functions, defined as per Equation 1, to give a single summary statistic. With this definition, the target value for our summary statistic was zero. We used a uniform perturbation kernel, N=5000 particles, and a schedule that reduced the rejection constant by 10% for each successive particle population. Further details on our implementation of the ABC-SMC algorithm can be found in the Methods section.

To assess the reproducibility of our parameter inference strategy, we performed two independent ABC trials that differed in their randomly generated initial particle population. We tested reproducibility by performing two-sample Kolmogorov-Smirnov hypothesis tests on the paired marginal distributions obtained in the two trials (α=0.05). The two trials exhibited no statistically significant difference in the distributions inferred for any parameter (p-values ranged from [0.218,0.758]). Thus, we concluded that the results were representative and reproducible. Figure 6 depicts the inferred marginal distributions for each model parameter, and Table 3 reports the corresponding 95% credible intervals (CI).

**Table 3.** Ninety-five percent credible intervals for plausible parameter values, as inferred via approximate Bayesian computation. Values are reported as log_10_(θ).

| parameter | 95% credible interval |
| --- | --- |
| $K_1$ | [-0.38, 3.9] |
| $K_3$ | [0.25, 3.9] |
| $K_4$ | [-3.2, 3.8] |
| $\alpha (= K_5/K_4)$ | [2.1, 4.0] |
| $K_6$ | [-3.9, 2.0] |
| $K_7$ | [-1.9, 3.9] |
| $\beta (= K_8/K_9)$ | [-2.7, -0.51] |
| $K_9$ | [-3.9, 1.9] |

**Fig 6.**
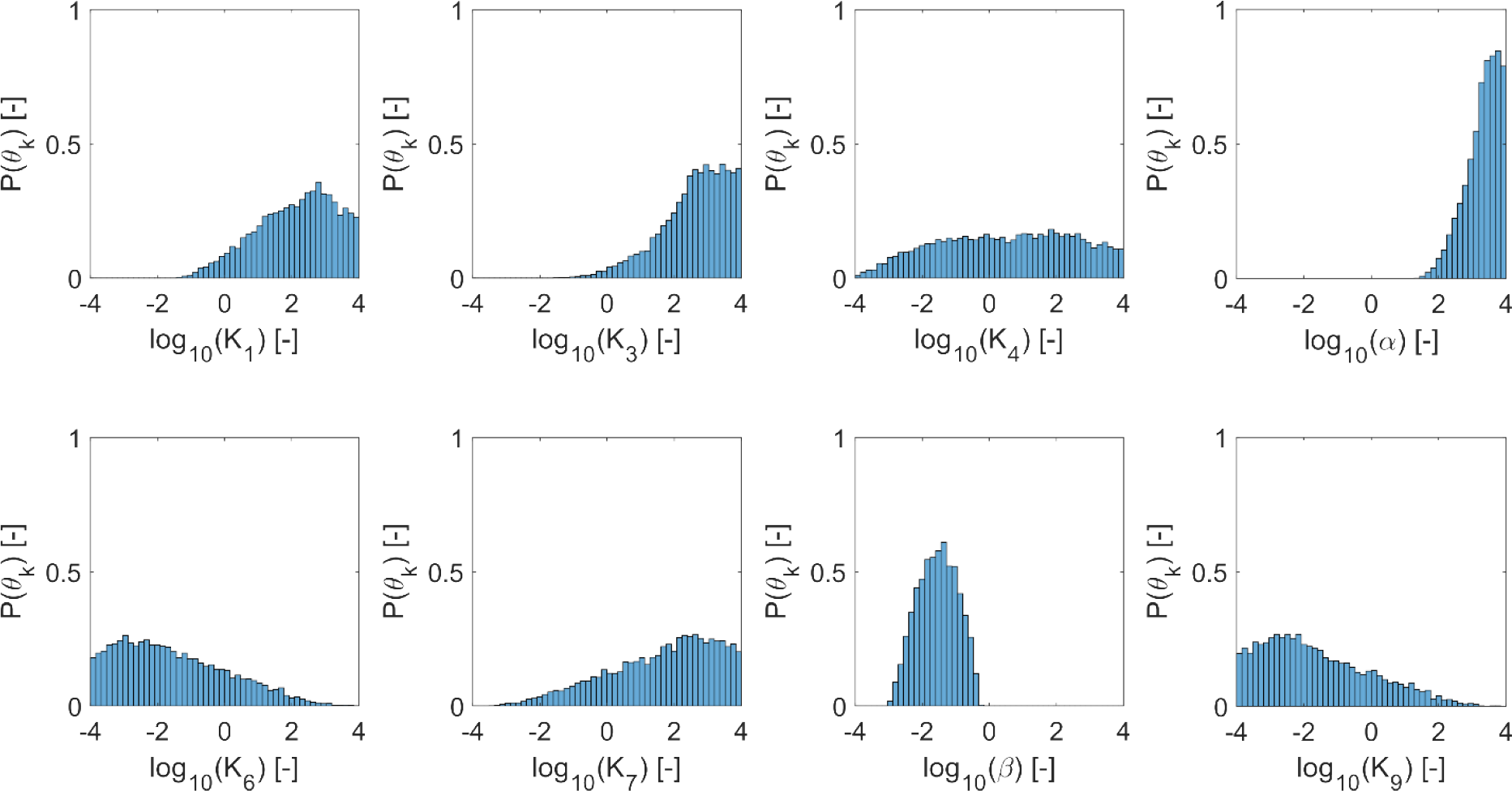
Marginal distributions indicating plausible parameter values, as inferred via approximate Bayesian computation. Inference was performed for two independent replicates using a particle population of 5000 parameter sets each. Plausible parameter sets are those that allow the model to satisfice all five criteria presented in Table 2 (see Table 1 for parameter descriptions). Satisficing these criteria implies that the model recapitulates the experimental observations of interest. The plots represent the final particle populations from both replicates, yielding histograms reflecting N=10000 satisficing parameter sets.

Although no parameters were well-constrained, our inference approach did identify plausible sub-domains in parameter space. Alternatively, one could say that we effectively excluded implausible domains that would not be worthy of further interrogation. Having searched over eight orders of magnitude, our ABC-SMC algorithm found plausible values ranging over 2-7 orders of magnitude (Table 3). On the more-constrained end of the scale, α (the relative rate of HOPS and SNARE reassociation) and β (the relative rate of HOPS displacement by Sec17) varied over 2-3 orders of magnitude. At the other extreme, inferred values for K_4_ (the HOPS:*trans*-SNARE dissociation rate), K_6_ (the Sec17 recruitment rate), K_7_ (rate of Sec17 turnover from the membrane), and K_9_ (the rate of Sec17 association with bare *trans*-SNARE complexes) extended across at least 6 orders of magnitude. Indeed, the marginal distribution for K_4_ exhibited plausible values across nearly the full range examined. Despite the broad range of values deemed plausible, we learned general emergent principles when we examined the relationships between parameter pairs by plotting 2D histograms.

### Model predicts that closed stoma exhibit stalled trans-SNARE fusion complexes

As indicated by the inferred values for α, our ABC-SMC process identified plausible parameter domains where HOPS and SNARE complexes reassociate with a rate constant greater than the HOPS:*trans-*SNARE super-complex dissociates. While we found plausible values for the HOPS:*trans*-SNARE dissociation rate, K_4_, throughout the interrogated parameter range, we consistently observed that the reassociation rate was 2-4 orders of magnitude faster (Table 3). Indeed, an associated finding was the predominance of protein super-complexes rather than HOPS absence or free SNARE proteins in the pre-fusion steady state. A survey of the model’s steady state across all 10,000 inferred parameter sets consistently returned the result that HOPS complexes are in HOPS:*trans*-SNARE super-complexes prior to fusion signaling (95% CI for free HOPS abundance relative to HOPS:*trans*-SNARE super-complexes: [0.952, 0.952]). Our simulations similarly predicted that SNARE proteins are almost exclusively in super-complexes rather than the free SNARE state (95% CI: [0.992, 1.00]). From this, we offer the testable hypothesis that HOPS should be observed in the vacuole membranes of closed stoma, and an appropriate biophysical experiment should provide evidence of membrane HOPS being associated with SNARE proteins.

The other key protein in our simulations is Sec17. We observed that the predicted abundance of Sec17 in the membrane is consistently lower than that of HOPS. Upon surveying the ratio of total membrane Sec17 to total HOPS across simulations based on our inferred parameter sets, we obtained steady-state values of 5.0e-5 (median; 95% CI: [2.2e-8,0.017]). Inspecting the plausible parameter sets indicates this may be due to differences in protein recruitment rates. When we examined the 2D histogram of HOPS and Sec17 turnover rates (K_1_ and K_7_, respectively), we found that turnover rates for the proteins may be comparable (Figure 7A). If not comparable, either protein might exhibit the greater rate of turnover. However, the relationship between rates of recruitment was less ambiguous. Two-D histograms suggest that HOPS and Sec17 recruitment rates (K_1_ and K_6_, respectively) may differ considerably (Figure 7B). Indeed, the modal outcome corresponds to HOPS being recruited at rates ∼5 orders of magnitude faster than Sec17. With Figure 7C, we introduce the additional observation that Sec17 turnover is typically faster than Sec17 recruitment (Figure 7C). From these observations, we offer the testable hypothesis that Sec17 should not be readily observed in the vacuole membranes of closed stoma. However, not seeing the protein under these conditions should not be misconstrued as implying that Sec17 could not have a role in vacuole membrane fusion.

**Fig 7.**
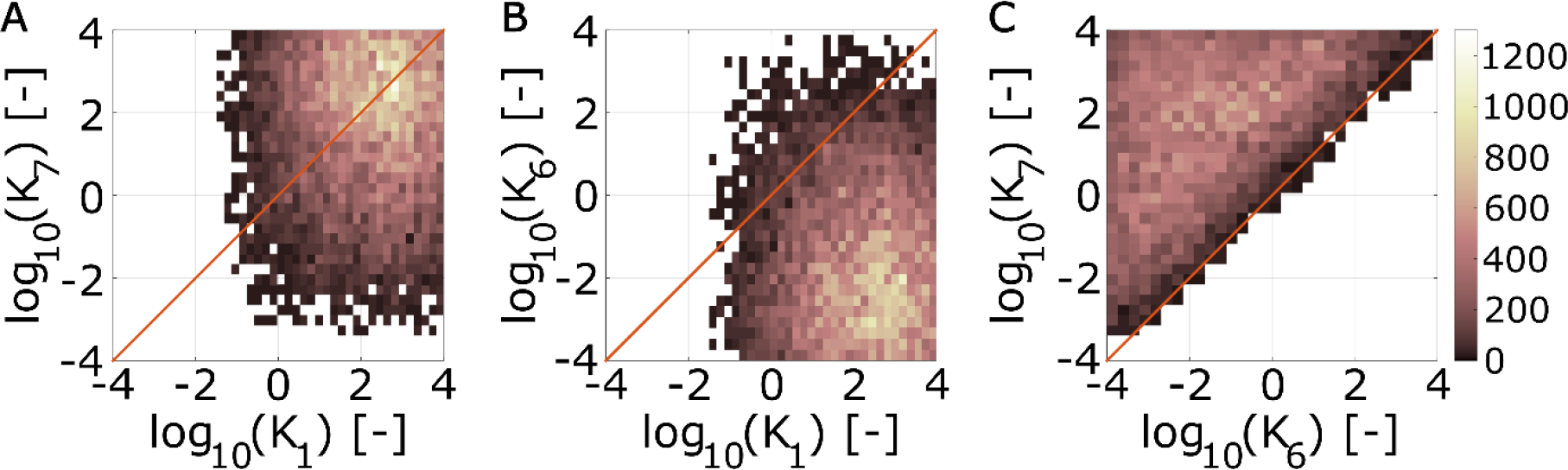
Plausible parameter domains indicate distinguishing recruitment rates for membrane HOPS and Sec17. Two-dimensional histograms depicting parameter values that satisfice all simulation criteria. (A) HOPS and Sec17 turnover rate constants (K_1_ and K_7_, respectively). (B) HOPS and Sec17 recruitment rate constants (K_1_ and K_6_, respectively). (C) Rate constants for Sec17 recruitment and turnover (K_6_ and K_7_, respectively). Plots represent combined parameter sets from two ABC trials, giving a total of N=10^4^ parameter sets satisficing all simulation criteria (see Table 2).

Lastly, our model frames the guard cell treatments that induce fusion as doing so via two different paths: one that proceeds by Sec17 engaging spontaneously with *trans*-SNARE complexes, and one that proceeds via Sec17 actively displacing HOPS from the HOPS:*trans*-SNARE super-complex (via events 8 and 9, respectively, in Figure 4). Via our inference algorithm, we estimated β – the ratio between the rate constants governing these paths (K_8_/K_9_). We observed this value to be consistently negative in log space (median -1.5; 95% CI [-2.7, -0.51]). The inferred range of values implies that the process requiring HOPS displacement proceeds at less than half the rate that it would if the *trans*-SNARE complex were not associated with HOPS (median ratio of 0.029; 95% CI: [0.0022, 0.31]). This suggests that Sec17 association with *trans*-SNARE complexes is hindered by the presence of HOPS.

### Modeling positions HOPS as a dual regulator of guard cell vacuole fusion, encouraging formation of the trans-SNARE fusion complex, but stalling the complex’s activity

By carefully integrating our knowledge of the molecular machinery involved in vacuole fusion with our observations of emergent vacuole morphology, we have arrived at a novel hypothesis (Figure 8) regarding the function of HOPS in plant guard cells. We predict that HOPS complexes promote the formation of a stable HOPS:*trans*-SNARE super-complex, but that super-complex cannot facilitate fusion until an appropriate signal is perceived. We thus frame a regulatory role for HOPS that is distinct from its chaperoning activity. Chaperoning helps form the fusion machinery, but our model suggests that this event is decoupled from fusion activity. In fact, analysis of the model indicates that the presence of HOPS hinders fusion activity. Furthermore, a sensitivity analysis constrained to the model’s plausible parameter domains indicates that the fusion rate may be insensitive to the chaperoning rate. We thus posit that a biological signal capable of triggering fusion should ultimately impact the HOPS:*trans*-SNARE super-complex, and it should do so by facilitating HOPS displacement. This impact could arise via changes in HOPS/SNARE binding affinity or Sec17/SNARE interactions. Such a change could, in turn, be induced by post-translational modification of a relevant protein – *e*.*g*., a HOPS subunit. Whether this change is mediated by phosphorylation status, an allosteric regulatory interaction, or another mechanism as yet unknown remains a topic for future investigation.

**Fig 8.**
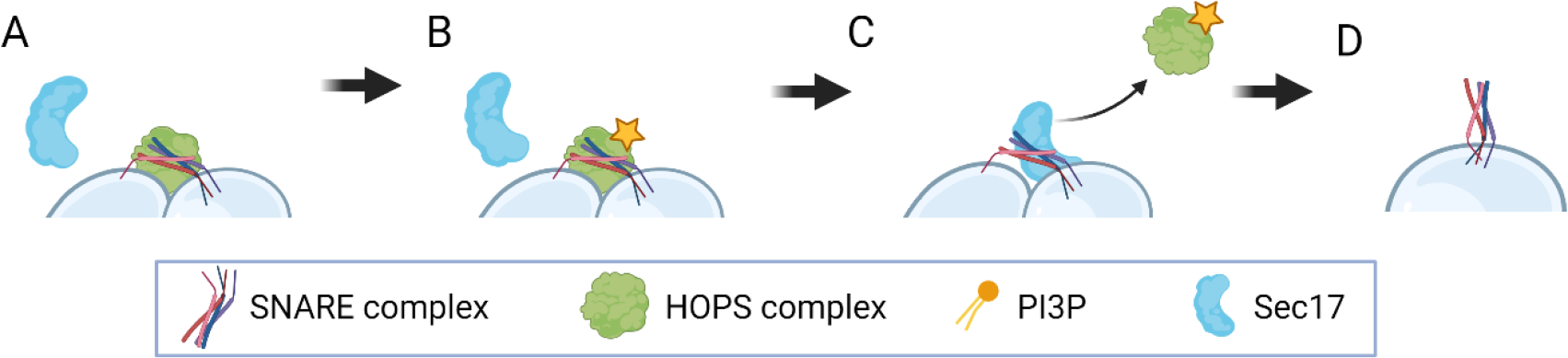
Modeling positions HOPS as a dual regulator of guard cell vacuole fusion. (A) HOPS promotes formation of the *trans*-SNARE fusion machinery, but then prevents fusion activity by hindering the access of Sec17. (B) A biological signal capable of reducing HOPS:*trans*-SNARE binding affinity (*e*.*g*., a signal resulting in the post-translational modification of a HOPS subunit, as indicated by the yellow star) could create the conditions required for Sec17 to displace HOPS and (C) thereby activate the otherwise stalled fusion complex, resulting in (D) membrane rearrangement and vacuole fusion.

Finally, we note that the proposed scheme positions the signal for vacuole fusion as one acting downstream of forming the *trans*-SNARE fusion complex. For example, in the plant-specific phenomenon of stomatal response to daylight, this mechanism would allow the plant to pre-dock pairs of fragmented vacuoles by forming stalled *trans*-SNARE complexes overnight. The system would then be poised to respond rapidly at daylight by producing a burst of fusion activity to support guard cell swelling and stomatal opening.

## Conclusion

We began this study with the unexpected biological observation that chemical depletion of a specific phosphoinositide – one apparently required to assemble fusion machinery in guard cells – induces fusion. Using simulation-based inference, we integrated existing biological knowledge with scarce phenotypic data to establish plausible kinetics associated with a candidate systems model. As data to inform the model, we considered the observed state of vacuole fragmentation in live cells under two lab-based perturbations: (i) a fusicoccin treatment that mimics the normal cues for stoma opening and (2) a wortmannin treatment that depletes the regulatory lipid of interest, PI3P. Taking this phenotypic data to be evidence of relative fusion rates, we implemented a model that predicts fusion activity emerging from a multi-step signaling pathway. Our observations regarding the state of fragmentation were few and qualitative, but they proved sufficient information to constrain a search for plausible model parameters.

Using an ODE model to make forward predictions and a Bayesian inference approach to reverse engineer governing parameters, we characterized those regions in the model’s multidimensional parameter space that are consistent with the expected fusion dynamics. Then, by sampling from that domain of plausible kinetics, we generated falsifiable mechanistic hypotheses regarding the intracellular localization of HOPS and SNARE complexes prior to fusion. We predicted that the apparently contradictory observation that PI3P is required for fusion, but removing PI3P promotes spontaneous fusion in plants, can be resolved by positing that HOPS and SNARE proteins exist as pre-formed, but stalled, HOPS:*trans-*SNARE super-complexes in the guard cells of closed stoma. Our work thus positions HOPS as having a dual role in regulating *trans-*SNARE fusion complexes in *Arabidopsis* guard cells. Namely, HOPS acts as both a promoter and inhibitor of vacuole fusion, executing these roles at different points in the non-linear signaling pathway that regulates vacuole fusion. We propose that the HOPS complex chaperones individual SNARE proteins into their *trans-* SNARE fusion machinery and then acts as a brake on the function of that machinery. Environmental cues for stoma opening would then act as a signal to release that brake.

Our model also introduces a functional role for a protein acting to displace HOPS from the *trans*-SNARE complex. This protein interacts with *trans*-SNARE assemblies to form a fusion-competent complex capable of rearranging membranes. By leveraging information from other kingdoms of life, we inferred that the yeast protein Sec17 could serve this function. Furthermore, apparent homologs of Sec17 (ASNAP1 and ASNAP2) exist in *Arabidopsis*. Additionally, ASNAP2 has been shown to interact with the vacuolar Qa SNARE SYP22 in root tissue isolates [40]. We suggest that research efforts focused on ASNAP1 and ASNAP2 would be useful avenues for future investigation. Whether ASNAP proteins interact with the specific SNARE hetero-tetramer involved in guard cell vacuole fusion remains unknown.

In addition to the insights documented here, our proposed model for vacuole fusion dynamics lays the foundation for future research as a tool for hypothesis generation. This will facilitate study of protein functions and interactions that would otherwise be difficult to track experimentally. Finally, future validation of this model may eventually lead to genetic applications to increase water use efficiency in dicot crop systems.

## Methods

### Plant growth conditions and stomata assays

Wild type *Arabidopsis thaliana* ecotype Columbia-0 (Col-0) plants were grown in soil at 22°C with a 16 h photoperiod. Leaves from 4-week old plants were cut in the morning and immediately processed to generate epidermal peels as described [41–43] with modifications. Briefly, a small leaf fragment was applied abaxially to a coverslip coated with medical adhesive and all but the bottommost layers of abaxial cells containing the stomata were scrapped away with a razor. 1 mm thick silicone isolators (GraceBio #664170) were used to create wells around the adhesive for incubations. Epidermal peels were immediately incubated in stomata buffer (MES pH 6.1). To induce stomatal closure, peels were incubated in closing buffer (10 mM MES pH 6.1, 40 mM malate, 5 mM CaCl_2_, 10 μM 2’,7’-Bis-(2-CarboxyEthyl)-5-(and-6)-CarboxyFluorescein, Acetoxymethyl Ester (BCECF, Fisher Scientific B1170), 50 mM abscisic acid (ABA, Sigma Aldrich A1049) at 22°C in the dark for two hours. To induce stomatal opening or vacuole fusion, ABA-treated peels were incubated in opening buffer (10 mM MES pH 6.1, 50 mM KCl) supplemented with 10 μM BCECF and either 3 μM fusicoccin (Sigma F0537) or 33 μM wortmannin (Sigma W3144). Wortmannin-treated peels were kept in the dark, while fusicoccin-treated peels were exposed to the light for up to 1 h. All concentrated stocks were first dissolved in DMSO. Images of leaf epidermal peels were acquired after ABA incubation and after 20, 40 and 60 min of fusicoccin or wortmannin treatments.

### Microscopy

Confocal laser scanning microscopy was carried out in a Zeiss LSM 980 confocal microscope. Images were taken with a 40× FCS water objective (1.1 N.A.). Acquisition of BCECF fluorescence was accomplished with 405 nm laser excitation and 495-550 nm emission filter set. Images were acquired with an Airyscan detector with a pinhole size of 2.5 airy units.

### Protein sequence comparisons

Sequence comparisons were performed using BLASTP 2.9.0+ comparing the Araport 11 protein dataset and the yeast protein sequence for Sec17 taken from the *Saccharomyces* Genome Database (SGD) [44–48].

### Governing equations of the mathematical model

Our model of consists of the following set of ordinary differential equations:

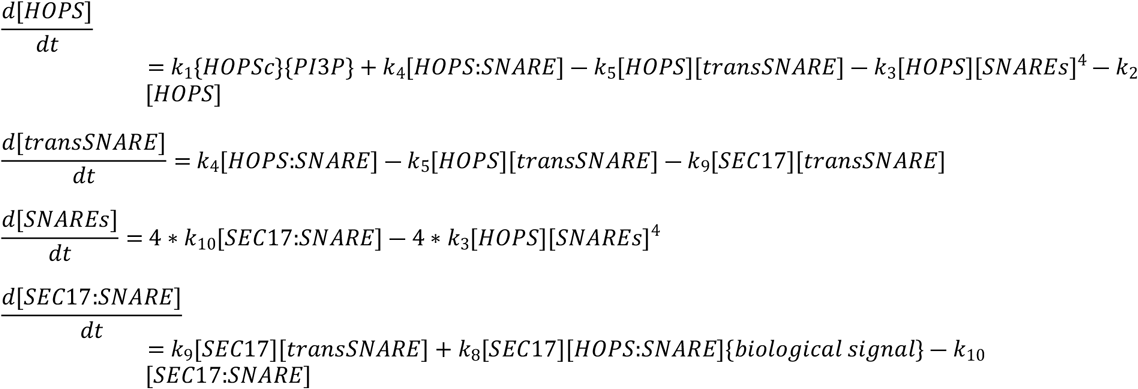

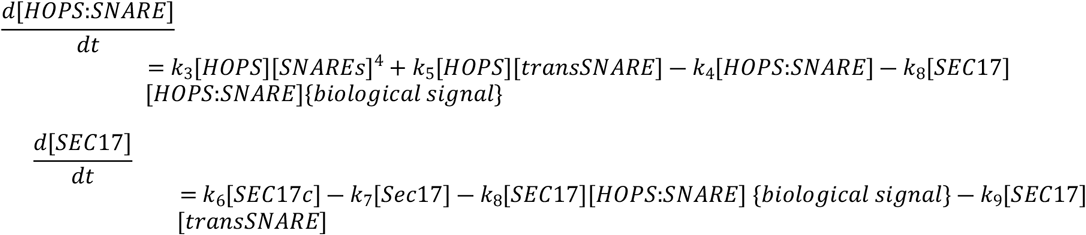

The quantities enclosed in curly brackets are Boolean variables indicating whether the indicated species is present or absent. We abstracted the availability of cytosolic HOPS subunits into a single cytosolic species that we denote as HOPSc. We assume that VPS41 recruitment acts as the rate-limiting step for recruitment of all HOPS subunits to form the HOPS complex at the membrane VPS41 is the subunit whose recruitment has been observed to be regulated by PI3P. Treating the HOPS subunits as abundant in the cytosol (*i*.*e*., with concentration not meaningfully altered by membrane recruitment) allowed us to fold the cytosolic abundance of HOPS into the rate constant for HOPS recruitment. We then treated the presence of cytosolic HOPS as binary variable {HOPSc}.

We non-dimensionalized the system of ODEs using a concentration scale of k_1_/k_2_ (*i*.*e*., the ratio of the rate constants for HOPS recruitment from the cytoplasm and HOPS turnover from the membrane) and a time scale of 1/k_10_ (*i*.*e*., the inverse of the rate constant for fusion). The dimensionless rate constants and their constituent parameters are listed in Table 4. This scaling reduced the number of model parameters from nine to eight. It also allowed us to express all rate constants relative to that for the fusion event, which has a non-dimensional value of one. In our analyses, for all parameters in our dimensionless system of equations, we considered rate constants four orders of magnitude larger and four orders of magnitude smaller than this reference value of one.

**Table 4.** Dimensionless parameters and their definitions.

| event | dimensionless rate constant | definition |
| --- | --- | --- |
| 1 | $K_2$ | $k_2/k_{10}$ |
| 2 | $K_2$ | $k_2/k_{10}$ |
| 3 | $K_3$ | $k_3k_1^4/(k_{10}k_2^4)$ |
| 4 | $K_4$ | $k_4/k_{10}$ |
| 5 | $K_5$ | $k_1k_5/k_2k_{10}$ |
| 6 | $K_6$ | $k_6/k_{10}$ |
| 7 | $K_7$ | $k_7/k_{10}$ |
| 8 | $K_8$ | $k_1k_8/k_2k_{10}$ |
| 9 | $K_9$ | $k_1k_9/k_2k_{10}$ |
| 10 | $K_{10}$ | 1 |

### Sobol’ global sensitivity analysis

The Sobol’ algorithm provides a point estimate of the Sobol’ index associated with each parameter and outcome. However, efficient calculation of the Sobol’ indices requires a numerical approximation involving many samples across the interrogated parameter domain. To determine whether each estimated index could credibly be differentiated from zero (at the chosen level of sampling), we introduced a dummy parameter and a dummy outcome to the analysis and used them as a control for true insensitivity. The dummy parameter does not feature in the systems model and thus provides a negative control for sensitivity of model outcomes. We set the dummy outcome equal to one of the model kinetic parameters, so that the dummy outcome provides an example of (i) perfect variance (T_j_ = 1) for that outcome-parameter combination and (ii) true insensitivity for all other parameters. We then used a Wilcoxon ranked sum test to determine whether, at a given sampling level, we could reject the null hypothesis that the distribution of bootstrapped sensitivity values for any given outcome-parameter pair has a median matching the negative control. We interpreted rejection of the null hypothesis (at a confidence level of p < 0.05) as evidence of a statistically significant sensitivity index.

### Parameter estimation using ABC-SMC

We performed parameter estimation using the approximate Bayesian computation with sequential Monte Carlo (ABC-SMC) approach described by Toni *et al*. [39]. We began with an uninformative uniform prior (defined in log space) for every model parameter and used uniform Sobol’ sampling to generate an initial population of N=5000 particles. We then solved the system of ODEs and evaluated the summary statistic for each particle. As per the ABC-SMC algorithm, we randomly selected a particle for perturbation – initially using uniform weighting. The particle’s parameters were then perturbed using a Markov kernel. We chose a uniform proposal kernel, 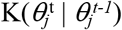, defined as per Equation 2, where*θ*_*j*_ denotes the value of parameter *j, t-1* denotes the current particle population, and *t* denotes the subsequent population. This limited the perturbation range for parameter *j* to within a distance *D* of the parameter’s current value. We defined *D* as 0.25Δ_*j*_, where Δ_*j*_ is the range of *θ*_*j*_ values represented in the current particle population.

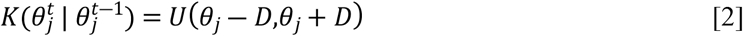

We rejected the proposed particle perturbation if the summary statistic evaluated for the perturbed parameter values exceeded a rejection constant, ε. Initially, we set the rejection constant to the 99^th^ percentile of the summary statistics characterizing the particle population. We then reduced ε by 10% for each subsequent population (*i*.*e*., ε_t_ = 0.9ε_t-1_). If the perturbed particle was rejected, we returned to the particle sampling step and repeated the process of particle selection, perturbation, and evaluation until an acceptable perturbation was identified. We then assigned that accepted particle to the next particle population and iterated until we identified a complete set of N new particles. Subsequent rounds of particle perturbation employed importance sampling to weight the selection of candidate particles to perturb. By iteratively reducing the rejection constant, the algorithm moved each subsequent population closer to the target distribution we sought – *i*.*e*., one that reflects a domain in parameter space plausibly describing our biological system. We iteratively generated new populations until 99% of the particles reflected a summary statistic of zero, where zero indicates a simulation that satisfies all acceptance criteria.

### Programming languages and code availability

The codes for this model and associated analyses were written in MATLAB (R2022B) (Inc. n.d.). Data and code used to generate the figures and perform the analyses for this paper are hosted at https://gitlab.com/hodgenscode/hodgens2023. A copy of the microscopy data has been made available at Zenodo under DOI:10.5281/zenodo.8408018.

## Acknowledgements

This work was supported by the National Science Foundation (MCB-1918746 award to M.R.P. and B.S.A.). Co-author DT Flaherty acknowledges fellowship support from NIH 5T32GM133366 (PIs Robert M. Kelly and Jason Haugh). We also thank Xiaohan Yang for help reviewing this manuscript. Figure 1 and Figure 8 were created using BioRender.com.

## Author contributions (CRediT taxonomy)

Investigation: CH, BSA, MRP, DTF, IK, AMP, LMG, NJE

Conceptualization: BSA, MRP

Funding Acquisition: BSA, MRP

Writing-original draft: CH, BSA

Writing-review & editing: BSA, CH, DTF, AMP, MRP

Methodology: CH, BSA, DTF

Software: CH, DTF

Formal Analysis: CH, DTF

## Supporting Information

Supplemental Table 1. Guard cell vacuole fragmentation status. Images of BCECF-stained guard cell vacuoles were qualitatively assessed to determine their fragmentation status. Individual guard cells were annotated as either fragmented (“F”), unfragmented (“U”), intermediate (“M”), or un-callable (“N”). Each row provides the number of guard cells for a given combination of treatment condition, acquisition date, treatment time, peel number, and fragmentation status. Source images used for this analysis are available on Zenodo under DOI:10.5281/zenodo.8408018.

## Data Availability Statement

Code used to perform the analyses for this paper are hosted at https://gitlab.com/hodgenscode/hodgens2023. Microscopy data used to generate the figures has been made available at Zenodo under DOI:10.5281/zenodo.8408018. Analyses were performed using MATLAB (R2022B).

## Financial Disclosure Statement

This work was supported by the National Science Foundation (MCB-1918746 award to M.R.P. and B.S.A.) The funders had no role in study design, data collection and analysis, decision to publish, or preparation of the manuscript.

## Competing interests

The authors have no competing interests to declare.

## Notes

### Competing Interest Statement

The authors have declared no competing interest.

